# Novel pedigree analysis implicates DNA repair and chromatin remodeling in Multiple Myeloma risk

**DOI:** 10.1101/137000

**Authors:** Rosalie G. Waller, Todd M. Darlington, Xiaomu Wei, Michael J. Madsen, Alun Thomas, Karen Curtin, Hilary Coon, Venkatesh Rajamanickam, Justin Musinsky, David Jayabalan, Djordje Atanackovic, Vincent Rajkumar, Shaji Kumar, Susan Slager, Mridu Middha, Perrine Galia, Delphine Demangel, Mohamed Salama, Vijai Joseph, James McKay, Kenneth Offit, Robert J. Klein, Steven M. Lipkin, Charles Dumontet, Celine M. Vachon, Nicola J. Camp

## Abstract

The high-risk pedigree (HRP) design is an established strategy to discover rare, highly-penetrant, Mendelian-like causal variants. Its success, however, in complex traits has been modest, largely due to challenges of genetic heterogeneity and complex inheritance models. We describe a HRP strategy that addresses intra-familial heterogeneity, and identifies inherited segments important for mapping regulatory risk. We apply this new Shared Genomic Segment (SGS) method in 11 extended, Utah, multiple myeloma (MM) HRPs, and subsequent exome sequencing in SGS regions of interest in 1063 MM / MGUS (monoclonal gammopathy of undetermined significance – a precursor to MM) cases and 964 controls from a jointly-called collaborative resource, including cases from the initial 11 HRPs. One genome-wide significant 1.8 Mb shared segment was found at 6q16. Exome sequencing in this region revealed predicted deleterious variants in *USP45* (p.Gln691*, p.Gln621Glu), a gene known to influence DNA repair through endonuclease regulation. Additionally, a 1.2 Mb segment at 1p36.11 is inherited in two Utah HRPs, with coding variants identified in *ARID1A* (p.Ser90Gly, p.Met890Val), a key gene in the SWI/SNF chromatin remodeling complex. Our results provide compelling statistical and genetic evidence for segregating risk variants for MM. In addition, we demonstrate a novel strategy to use large HRPs for risk-variant discovery more generally in complex traits.

**AUTHOR SUMMARY:** Although family-based studies demonstrate inherited variants play a role in many common and complex diseases, finding the genes responsible remains a challenge. High-risk pedigrees, or families with more disease than expected by chance, have been helpful in the discovery of variants responsible for less complex diseases, but have not reached their potential in complex diseases. Here, we describe a method to utilize high-risk pedigrees to discover risk-genes in complex diseases. Our method is appropriate for complex diseases because it allows for genetic-heterogeneity, or multiple causes of disease, within a pedigree. This method allows us to identify shared segments that likely harbor disease-causing variants in a family. We apply our method in Multiple Myeloma, a heritable and complex cancer of plasma cells. We identified two genes *USP45* and *ARID1A* that fall within shared segments with compelling statistical evidence. Exome sequencing of these genes revealed likely-damaging variants inherited in Myeloma high-risk families, suggesting these genes likely play a role in development of Myeloma. Our Myeloma findings demonstrate our high-risk pedigree method can identify genetic regions of interest in large high-risk pedigrees that are also relevant to smaller nuclear families and overall disease risk. In sum, we offer a strategy, applicable across phenotypes, to revitalize high-risk pedigrees in the discovery of the genetic basis of common and complex disease.

## Introduction

Rare risk variants have been suggested as a source of missing heritability in the majority of complex traits [1–3]. High-risk pedigrees (HRPs) are a mainstay for identifying rare, highly penetrant, Mendelian-like causal variants [4–11]. However, while successful for relatively simple traits, genetic heterogeneity remains a major obstacle that reduces the effectiveness of HRPs for gene mapping in complex traits [12,13]. Also challenging is mapping regulatory variants, likely to be important for complex traits, necessitating interrogation outside the well-annotated coding regions of the genome [14,15]. Localizing chromosomal regions to target the search for rare risk variants will be instrumental in mapping them.

Here we develop a HRP strategy based on our previous Shared Genomic Segment (SGS) approach [16] that focuses on pedigrees sufficiently large to singularly identify segregating chromosomal segments of statistical merit. The method addresses genetic heterogeneity by optimizing over all possible subsets of studied cases in a HRP. Key to the utility of the method is the derivation of significance thresholds for interpretation. These thresholds address the genome-wide search and the multiple testing, inherent from the optimization, through use of distribution fitting and the Theory of Large Deviations.

We apply this novel method to 11 MM HRPs, and use exome sequencing from a collaborative resource of 55 multiplex MM or MM/MGUS pedigrees to perform subsequent targeted searches at the variant level. MM is a complex cancer of the plasma cells with 30,330 new cases annually (incidence 6.5/100,000 per year) [17]. Despite survival dramatically increasing from 25.8% in 1980 to 48.5% in 2012, MM remains a cancer with one of the lowest 5-year survival rates in adult hematological malignancies [17]. MM is preceded by a condition referred to as monoclonal gammopathy of undetermined significance (MGUS). Evidence for the familial clustering of MM is consistently replicated [18–21], as is its clustering with MGUS [22–25]. Genetic pedigree studies in MM are scarce as it remains a challenge to acquire samples in pedigrees due to rarity and low survival rates. The Utah MM HRPs are one of only a few pedigree resources worldwide and contains unparalleled multi-generational high-risk pedigrees. Thus far, no segregating risk variants have been identified for MM.

## Results

### Pedigree analysis strategy

We developed a gene mapping strategy, based on the SGS method [16,26], that accounts for intra-familial heterogeneity and multiple testing. The basic SGS method identifies all genomic segments shared identical-by-state (sharing without regard to inheritance) between a defined set of cases using a dense genome-wide map of common single nucleotide polymorphisms (SNPs), either from a genotyping platform or extracted from sequence data. If the length of a shared segment is significantly longer than by chance, inherited sharing is implied; theoretically, chance inherited sharing in distant relatives is extremely improbable. Nominal chance occurrence (nominal p-value) for shared segments is assessed empirically using gene-drop simulations to create a null distribution, as follows. Null genotype configurations are generated by assigning haplotypes to pedigree founders according to a publicly available linkage disequilibrium (LD) map, followed by segregation of these through the pedigree structure to the case set via simulated Mendelian inheritance according to a genetic (recombination) map. Gene-drops are performed independent of disease status and the resulting genotype data in the case set are representative of chance sharing. This basic method was shown to have excellent power in homogeneous pedigrees [16].

In our new strategy, we iterate over all non-trivial combinations of the cases (subsets) in each pedigree to address heterogeneity in a “brute-force” fashion. For each subset, shared segments at every position throughout the genome are identified and nominal p-values assigned. Across subsets, an optimization procedure is performed at every marker across the genome to identify the segment with the most significant sharing evidence. All shared segments selected by the optimization procedure, and their respective p-values, comprise the final optimized SGS results.

To perform significance testing and identify segments that are unexpected by chance (hypothesized to harbor risk loci), we derive significance thresholds to account for the genomewide optimization. Acknowledging that the vast majority of observed sharing across a genome is under the null (true risk loci are a very small minority of the genome), we use the observed optimized results (*Y* = −*log*_10_(*p*), where *p* is the empirical p-value) to model the distribution for optimized SGS results. We note that this approach may be slightly conservative because signals for true risk loci are also included. We identified the gamma distribution as adequate to represent the distribution (Fig. 1). Based on the fitted distribution, *Y*~Γ(*k*,*σ*), where *k* and *σ* are the shape and rate parameters, we apply the Theory of Large Deviations; previously applied to successfully model genome-wide fluctuations in linkage analysis [27]. The significance threshold, *T*, accounts for multiple testing of optimized segments across the genome, and is found by solving Eq. 1:

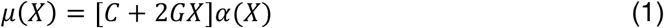

**Fig. 1.**
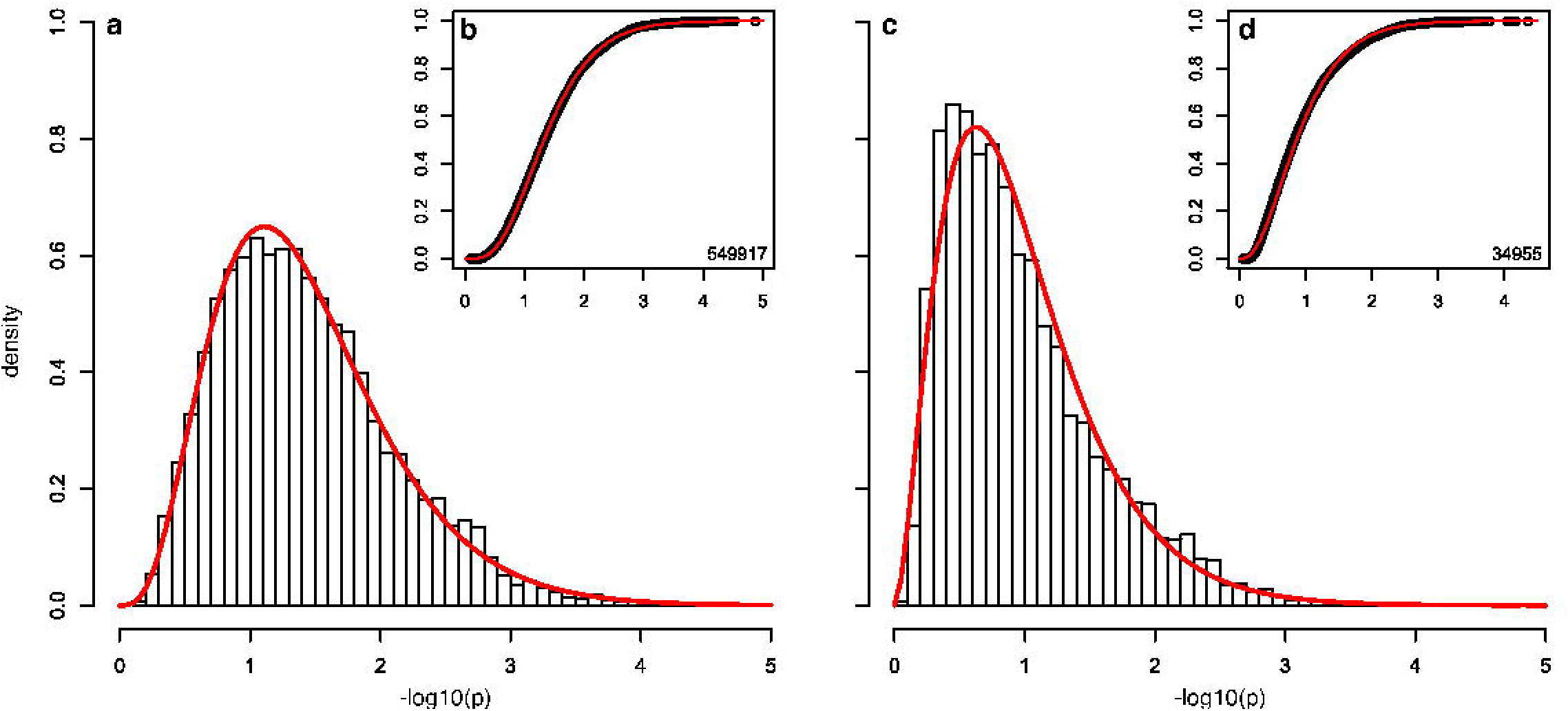
Adequacy of the gamma distribution. The gamma distribution provides an adequate fit for multiple types of pedigrees. For example, HRP 549917 has *k* = 4.4 and *σ* = 3.6 with good visual density (a) and CDF (b) fit, with = 0.9. (Goodness of fit was estimated with, the median of empirical chi-squared distribution divided by the median of the expected chi-squared distribution.) HRP 34955 has *k* = 2.8 and *σ* = 2.9 with good visual density (c) and CDF (d) fit, with *λ* = 1.0.

where, 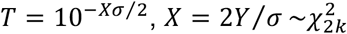 μ(*X*) is the genome-wide false positive rate required, is the number of chromosomes, *α*(*X*) is nominal probability of exceeding *X*,*G* and is the genome length in Morgans. A criterion of *μ*(*X*) = 0.05 is typically used to define the genome-wide significant threshold (false positive rate of 0.05 per genome), and *μ*(*X*) = 1 to define the genome-wide suggestive threshold (false positive rate of 1 per genome).

In general, we found that the fitted distributions produced stable significance thresholds after 100,000-300,000 simulations (Table 1). Typically, threshold determination requires 1,000-3,000 CPU hours per pedigree, increasing with the number of subsets and separating meioses between pedigree cases. For example, in pedigree UT-571744, 300k simulations genome-wide (2,513,408 segments) took 1,275 CPU hours on tangent nodes featuring Intel Xeon E5-2650 processors. Once significance thresholds are established, subset/segment combinations of potential interest are identified and additional simulations are restricted to those combinations to gain the required p-value resolution. For these subsequent targeted simulations, we use a marginalized LD map specific for the segment of interest, dramatically reducing the analysis time. For example, in pedigree UT-571744, 600M simulations on one segment took 325 CPU hours on tangent nodes featuring Intel Xeon E5-2650 processors. See S1 Fig. for an overview of the strategy pipeline.

**Table 1.** Genome-wide Significance Thresholds. Fitted distributions are stable enough for threshold determination after 100,000 to 300,000 simulations.

| Pedigree | 100k | 200k | 300k | 1M |
| --- | --- | --- | --- | --- |
| 260 | $6.36 \times 10^{-6}$ | $6.35 \times 10^{-6}$ | $6.28 \times 10^{-6}$ | $6.25 \times 10^{-6}$ |
| 576834 | $3.50 \times 10^{-6}$ | $3.53 \times 10^{-6}$ | $3.53 \times 10^{-6}$ | $3.51 \times 10^{-6}$ |
| 571744 | $3.80 \times 10^{-6}$ | $3.83 \times 10^{-6}$ | $3.75 \times 10^{-6}$ | $3.80 \times 10^{-6}$ |
| 34955 | $5.67 \times 10^{-6}$ | $5.60 \times 10^{-6}$ | $5.61 \times 10^{-6}$ | $5.61 \times 10^{-6}$ |

### Application to Utah, MM HRPs

We applied our new pedigree analysis strategy to 11 Utah MM HRPs using high-density OMNI Express SNP array genotype data. Each pedigree was selected to contain excess MM (4-37 MM total per pedigree), had 2-4 sampled MM cases with genotype data, and 8-23 meioses per pedigree between the sampled cases. After quality control, a consistent set of 678,447 SNPs were used for all SGS analyses. The total number of shared segments for each pedigree across all subsets ranged from 638,525 to 6,765,500 (larger pedigrees with more subsets producing larger numbers of segments). After optimization, *Y* = −*log*_10_(*p*) for 6,697 to 10,369 segments were fit to gamma distributions for each pedigree, and used to determine genomewide significant and suggestive thresholds (Eq. 1). The genome-wide significant thresholds ranged from 6.2×10^-5^ to 7.8×10^-7^ and genome-wide suggestive from 8.2×10^-4^ to 2.1×10^-5^ (S1 Table).

A genome-wide significant, 1.8 Mb shared segment (p = 3.3x10-6) was observed in pedigree UT-571744. All three genotyped MM cases, separated by 20 meioses, share the segment (Fig. 2a and Table 2). The segment is located at chromosome 6q16 (98.49-100.24 Mb; hg19) and includes 9 genes: *POU3F2*, *FBXL4*, *FAXC*, *COQ3*, *PNISR*, *USP45*, *TSTD3*, *CCNC*, and *PRDM13* (Figure 2b).

**Fig. 2.**
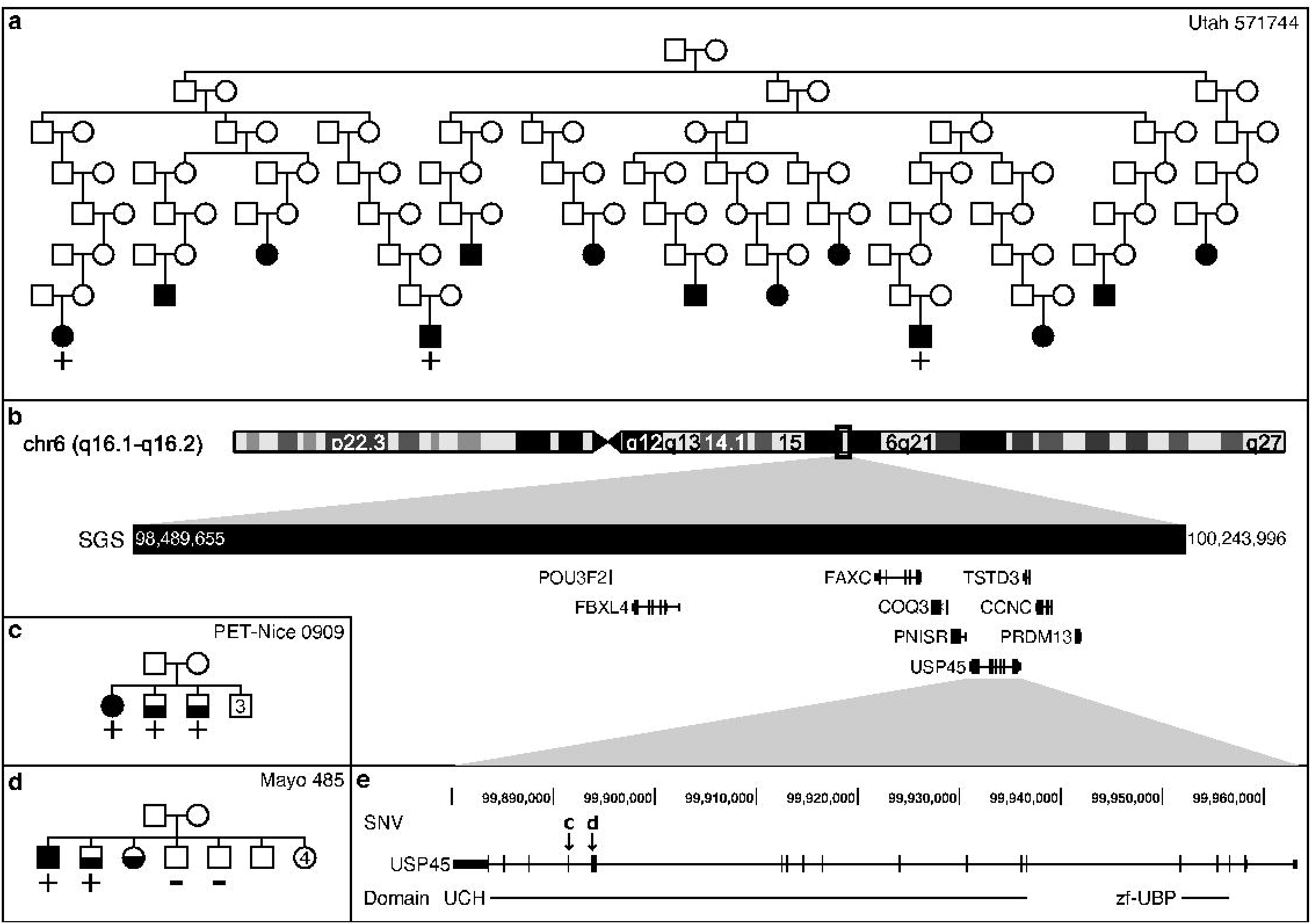
Significant SGS, pedigrees, and segregating SNVs. In pedigrees, MM cases are fully shaded and MGUS cases are half shaded. Numbers indicate multiple individuals. a) Utah pedigree, 571744, sharing the genome-wide significant SGS. The pedigree is trimmed to allow for viewing (37 MM confirmed cases are known in this pedigree, 3 were ascertained and genotyped). + indicates the genotyped MM cases that are SGS carriers, - indicates genotyped and non-carriers, no carrier status indicates not genotyped. Note - the genealogy extends beyond SEER cancer registry data. MGUS are unknown in this pedigree. b) Genomic region of significant SGS. c) INSERM pedigree carrying the stop gain SNV marked by “c” in box e. 1 MM and 2 MGUSs carry the SNV. d) Mayo Clinic pedigree carrying the missense SNV marked by “d” in box e. 1 MM and 1 MGUS carry the SNV, but not 2 unaffected siblings. e) Risk candidate gene, *USP45*, has 2 segregating SNVs in the ubiquitin C-terminal hydrolase 2 (UCH) domain.

**Table 2.**
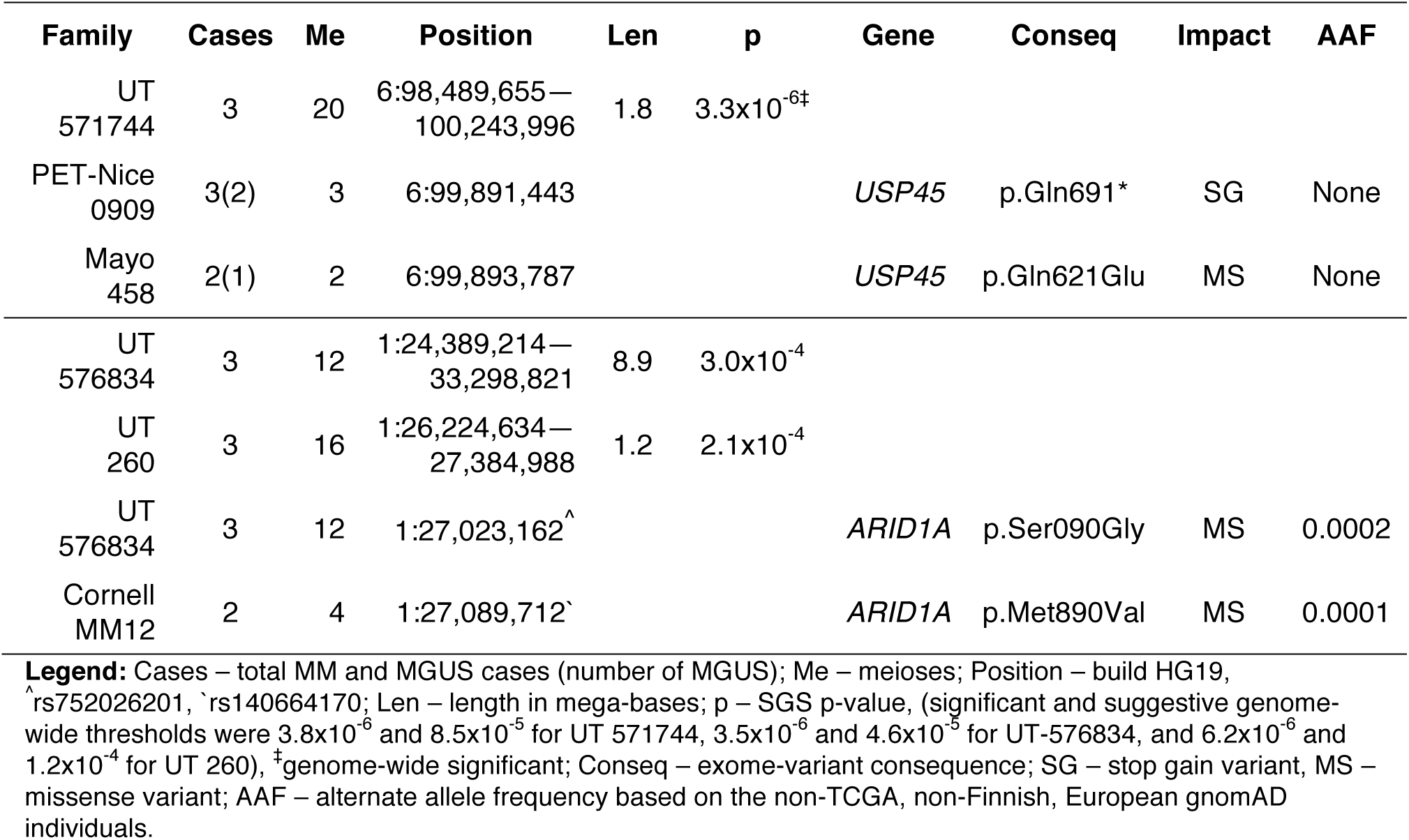
Significant or overlapping SGSs and segregating SNVs.

| Family | Cases | Me | Position | Len | p | Gene | Conseq | Impact | AAF |
| --- | --- | --- | --- | --- | --- | --- | --- | --- | --- |
| UT<br>571744 | 3 | 20 | 6:98,489,655—<br>100,243,996 | 1.8 | 3.3x10<sup>-6‡</sup> |  |  |  |  |
| PET-Nice<br>0909 | 3(2) | 3 | 6:99,891,443 |  |  | *USP45* | p.Gln691* | SG | None |
| Mayo<br>458 | 2(1) | 2 | 6:99,893,787 |  |  | *USP45* | p.Gln621Glu | MS | None |
| UT<br>576834 | 3 | 12 | 1:24,389,214—<br>33,298,821 | 8.9 | 3.0x10<sup>-4</sup> |  |  |  |  |
| UT<br>260 | 3 | 16 | 1:26,224,634—<br>27,384,988 | 1.2 | 2.1x10<sup>-4</sup> |  |  |  |  |
| UT<br>576834 | 3 | 12 | 1:27,023,162<sup>^</sup> |  |  | *ARID1A* | p.Ser090Gly | MS | 0.0002 |
| Cornell<br>MM12 | 2 | 4 | 1:27,089,712<sup>`</sup> |  |  | *ARID1A* | p.Met890Val | MS | 0.0001 |

Legend: Cases - total MM and MGUS cases (number of MGUS); Me - meioses; Position - build HG19, ^^^rs752026201, ^`^rs140664170; Len - length in mega-bases; p - SGS p-value, (significant and suggestive genome wide thresholds were 3.8x10^-6^ and 8.5x10^-5^ for UT 571744, 3.5x10^-6^ and 4.6x10^-5^ for UT-576834, and 6.2x10^-6^ and 1.2x10^-4^ for UT 260), ^‡^genome-wide significant; Conseq - exome-variant consequence; SG - stop gain variant, MS - missense variant; AAF - alternate allele frequency based on the non-TCGA, non-Finnish, European gnomAD individuals.

We also identified two HRPs, UT-576834 and UT-260, with overlapping shared segments at 1p36.11 (Fig. 3). A 8.9 Mb (24.39-33.30 Mb, p = 3.0×10-4) segment was observed in 3 of the 4 genotyped MM cases in UT-576834, shared across 12 meioses (Fig. 3b and Table 2). A nested 1.2 Mb shared segment (26.22-27.38 Mb; p = 2.1×10-4) segregated to 3 MM cases separated by 16 meioses in UT-260 (Fig. 3a and Table 2). The overlapping segment contains 30 genes (Fig. 3d).

**Fig. 3.**
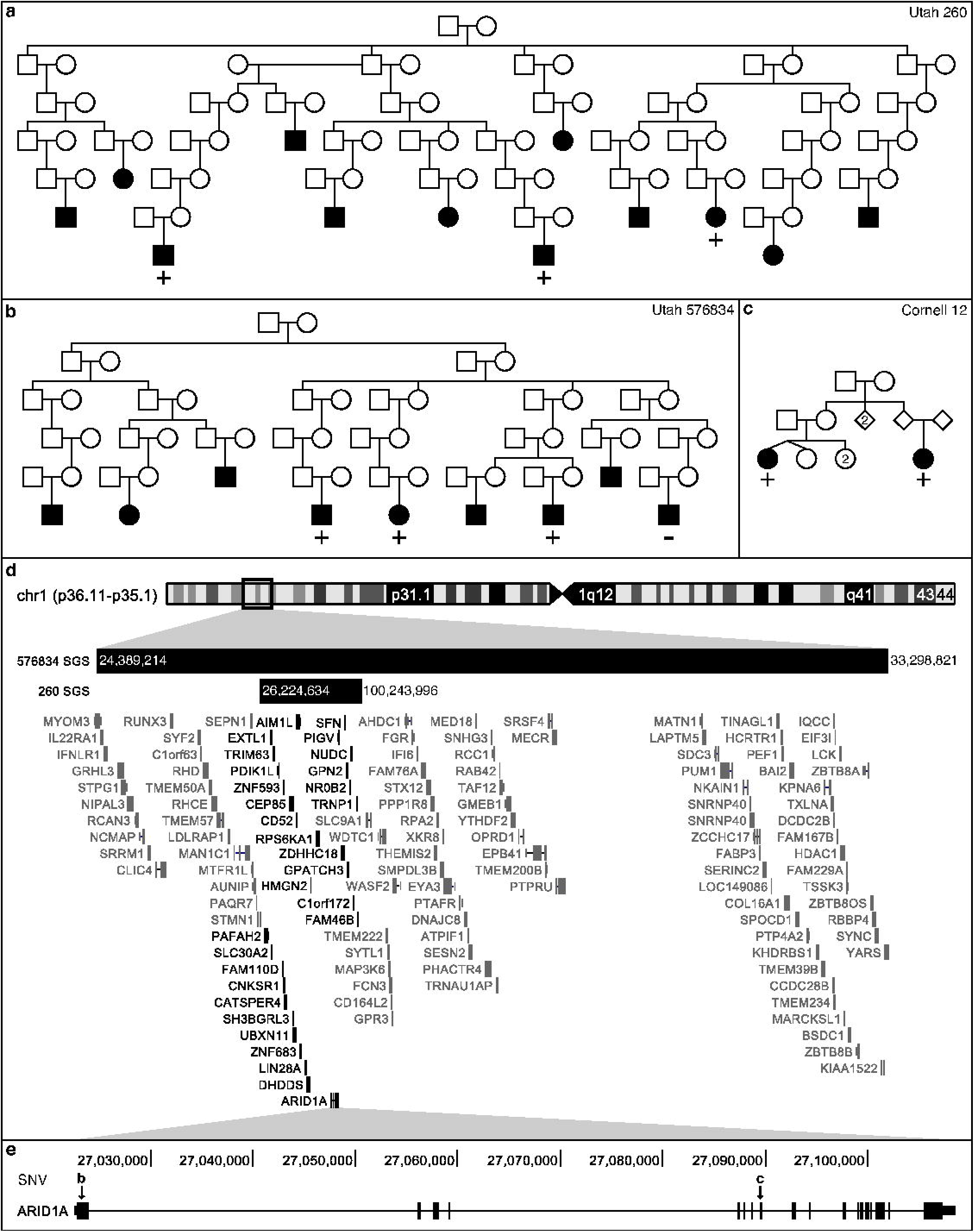
SGS with multiple lines of evidence. a/b) Utah pedigrees carrying the overlapping SGSs on chr1p36.11-p35.1. + indicates the genotyped MM cases that are SGS carriers, - indicates genotyped and non-carriers, no carrier status indicates not genotyped. c) Weill Cornell pedigree with a segregating, missense SNV in *ARID1A* indicated by “c” in e. d) Genomic region of overlapping SGS. Dark black genes fall in both regions. e) 2 rare and segregating, missense SNVs were observed in whole-exome sequencing. SNV “b” is carried by the cases indicated with + in box b. SNV “c” in carried by the cases in box c.

### Exome follow-up of shared segments in HRPs

Whole-exome sequencing (WES) data was interrogated, targeted to the identified SGS region, to identify potential risk variants in the pedigree sharers in the HRP and in a broader set of 44 pedigrees. WES data was available for: 28 cases from the 11 extended Utah HRPs; and 126 exomes from 44 densely clustered MM/MGUS families from Mayo Clinic Rochester, Weill Cornell, Memorial Sloan Kettering Cancer Center, International Agency for Research on Cancer, and INSERM France (S2 Table). Prioritization was used to identify variants that were: in the target segment; rare (alternate allele frequency, AAF<0.001 in the non-Finnish, European, gnomAD individuals), potentially deleterious (variant impact predicted to be high or moderate); and observed recurrently in the appropriate segment sharers (if observed in the segment discovery pedigree).

At 6q16, no rare, potentially deleterious coding risk variants were shared by the 3 UT-571744 MM cases in the 1.8 Mb genome-wide significant segment, indicating non-coding regulatory variants may be responsible for MM risk in this pedigree. However, two, rare coding and potentially deleterious single nucleotide variants (SNVs) were identified in two MM/MGUS families (Fig. 2c-e and Table 2). Both SNVs are in the hydrolase domain of *USP45*: a stop gain (p.Gln691*) shared by 3 sibling cases (1 MM and 2 MGUS) in an INSERM family (PET-Nice 0909) and a missense SNV (p.Gln621Glu) shared by 2 siblings (1 MM and 1 MGUS) but not their 2 screened unaffected siblings in Mayo family 485. Coverage of these positions in ExAC sequence data is high (> 99% of the 60,706 ExAC samples had at least 10x read coverage) and neither variant was observed. Collating the SGS evidence in UT 571744 (genome-wide rate of µ=0.0423) with the sequence findings, correcting for 11 SGS pedigrees, the 45 pedigrees interrogated for sequence variants, and the 9 genes in the SGS region, we estimate the rate of observing all these findings at the 6q16 region by chance is low (π=0.01, see Methods) and study-wide significant.

Pedigree exomes in the 1.2 Mb segment at 1p36.11 revealed two, rare and potentially deleterious SNVs. The first in discovery pedigree UT-576834: a missense SNV (rs752026201, p.Ser90Gly, AAF = 0.0002 in gnomAD) in *ARID1A* (Fig. 3e) shared by 3 of the 4 Utah MM cases, concordant with the segment sharing pattern. A second rare, missense SNV in *ARID1A* (rs140664170, p.Met890Val, AAF = 0.0001 in gnomAD) was found to be carried by a pair of MM cousins in Weill-Cornell family 12 (Fig. 3c and e, and Table 2). Based on the ExAC data, *ARID1A* is extremely intolerant to missense variants and loss of function (LoF) SNVs [28].

### Pathway follow-up of candidate genes

Our SGS findings and pedigree WES identify *USP45* and *ARID1A* as candidate genes for inherited MM risk. We further investigated shared segments and WES for evidence supporting the complexes USP45 and ARID1A are involved in. Here we further expanded our WES to: 186 MM/MGUS cases (early onset MM/MGUS or familial MGUS) from our collaborative group, 733 sporadic MM cases from dbGaP [29], and 964 controls [30].

USP45 is an essential DNA repair regulator, de-ubiquitylating ERCC1 to allow for DNA translocation of the ERCC1-ERCC4 endonuclease [31,32]. This endonuclease is a part of the global genome nucleotide-excision repair (GG-NER) incision complex, a 22 protein complex essential to removing lesions from DNA and cancer prevention [33–36] (S3 Table). We reviewed SGS results in the Utah HRPs at the location of these 22 genes and identified a genome-wide suggestive segment in pedigree UT-34955 (S2 Fig.). This HRP identified a 0.8 Mb segment at 19q13 (45.71-46.51 Mb; hg19), containing 31 genes including *ERCC1* and *ERCC2* (S2 Fig. and S4 Table). The segment is shared by 3 MM cases separated by 16 meioses (p = 6.6×10^-5^). No rare, coding variants were identified from the WES in the 3 MM cases in UT-34955, nor in the remaining 44 pedigrees/families. We interrogated the 23 GG-NER genes in our 919 MM/MGUS exomes. This identified a ClinVar-annotated pathogenic, missense SNV in *ERCC4* (p.Arg799Trp) in one early-onset MM case and one sporadic MM case, and a stop-gain SNV in *ERCC3* (p.Arg574Ter), in the same domain as a ClinVar-annotated pathogenic variant, in a second early-onset MM case (S4 Table). Further, burden testing in all MM cases vs controls was significant in 2 of the 23 GG-NER genes: *GTF2H1* and *DDB1* after correcting for multiple testing (S3 Table). The occurrence of two significantly burdened genes (at *α*=0.0022) from 23 genes is unexpected (p=0.0011, Binomial(23,0.0022)).

ARID1A is a member of the SWI/SNF chromatin remodeling complex, a 15 gene complex involved in DNA transcription regulation [37] (see S5 Table). Members of this complex are mutated in >20% of malignancies [38–40], but are extremely intolerant to LoF and missense variation [41] (S5 Table). We reviewed SGS results in the Utah HRPs at the location of these 15 genes and identified a marginal, genome-wide suggestive segment in pedigree UT-549917 shared by 4 MM cases across 21 meioses (p = 2.17×10-5, S3 Fig. and S6 Table). This 1.5 Mb segment at chr3p21.1-p21.2 (52.01-53.56 Mb; hg19) contains 32 genes including *PBRM1* from the SWI/SNF complex. No coding variants were identified in this gene in UT-549917, nor in the remaining 44 pedigrees/families. Burden testing was significant for 7 of the 15 genes in the complex after correcting for multiple testing: *ARID1A*, *ARID1B*, *SMARCA4*, *ACTL6A*, *SMARCD3*, *SMARCC2*, and *SMARCE1* (S5 Table). The occurrence of seven significantly burdened genes (at *α*=0.0033) from 15 genes is unexpected by chance (p=2.7×10^-14^, Binomial(15,0.0033)).

## Discussion

We developed a novel strategy to identify segregating chromosomal segments shared by subsets of cases in HRPs. It focuses on extended HRPs that are singularly powerful to identify significant genetic segregation. Our strategy allows for genetic heterogeneity within such pedigrees and provides formal significance thresholds for valid interpretation. Previously, extended HRP have not delivered on their potential in complex traits because in common, complex traits, HRPs are likely enriched for multiple susceptibility variants and may capture both familial and sporadic cases in their branches. Our optimization strategy over subsets is attractive because it allows for heterogeneity without prior knowledge of genetic similarities or deep phenotyping. This new statistic also identifies the sharers and clearly delimits the shared region, making follow-up interrogation straight-forward. This is a distinct advantage over standard linkage analysis and previous pairwise SGS methods where neither sharers or the region are defined [42].

Application of the method to extended MM pedigrees demonstrated the utility of this new method and illustrated that the segments identified were used successfully to narrow the search for risk variants in smaller pedigrees, allowing for an overall strategy that can utilize both large pedigrees and smaller families together for discovery (Table 2, Fig. 2 and Fig. 3). Post-hoc, additional value can be gained from demographic and/or clinical data on the sharing subsets shedding light on other shared characteristics that may aid future mapping. Also, we note that in the absence of any significant findings, genome-wide SGS results can be used as genomic annotations of segregation evidence for more heuristic approaches.

While we identified several rare, potentially deleterious coding variants of interest, several of the SGS discovery pedigrees had no coding variants that satisfied prioritization criteria. We believe this will be characteristic of complex traits and that regulatory variants will also play a substantial role. Mutations with strong causal likelihood found in other disease cohorts may focus the search for regulatory variation to particular genes within a shared segment, as with *USP45* in MM. In the absence of such compelling evidence, a return to pedigree segregation methods will provide identification of statistically compelling regions which can concentrate efforts to identify and characterize regulatory risk variants. Future work will include targeted sequencing of the promising MM SGS identified to investigate non-coding variants that may play a role in MM risk in these families. Our proposed method is a new analytic tool with the potential to reinvigorate the use of extended HRPs in the identification of risk variants that contribute to common, complex disease.

Multiple myeloma is a malignancy of the plasma cells that has been shown to be familial [43]. Consistent with a role for genetics, case-control studies have been successful in identifying association signals for 17 low-risk variants [44–48]. However, despite consistent evidence for familial clustering, our study is the first to explore high-risk MM pedigrees. Using the unique genealogical database available in Utah, we identified and studied extended MM HRPs. We identified a genome-wide significant segment containing *USP45*, an important regulator of DNA repair (Fig. 2 and Table 2), and a genome-wide suggestive segment harboring other genes in the GG-NER incision complex (*ERCC1* and *ERCC2*). Exome sequencing in a collaborative resource of high-risk families and early-onset cases revealed four rare, potentially deleterious coding variants; two novel variants in *USP45* segregating in two pedigrees and two variants in early-onset cases in *ERCC3* and *ERCC4*, the latter annotated as pathogenic in ClinVar. Burden testing including sporadic MM, and comparing to controls, identified significant enrichment for variants in MM cases in 2 of the 23 GG-NER genes in the protein endonuclease regulation complex.

In particular, the functional literature supports *USP45* as a candidate cancer risk gene. USP45 has been shown to deubiquitylate ERCC1, a catalytic subunit of the ERCC1-ERCC4 DNA repair endonuclease (ERCC4 also known as XPF) [31]. This endonuclease is a critical regulator of DNA repair processes [34]. The complex repairs recombination, double strand break, and inter-strand crosslink by cutting DNA overhangs around a lesion, degrades 3' G-rich overhangs in telomere maintenance, and plays a role in cancer prevention and in tumor resistance to chemotherapy [31,34]. Mouse models have shown USP45 knockout cells have higher levels of ubiquitylated ERCC1 and that cells are hypersensitive to UV radiation and DNA inter-strand cross-links, repair of UV-induced DNA damage, and ERCC1 translocation to DNA damage is impaired [31]. Hence, the deubiquitylase activity of USP45 is important for maintaining the DNA repair ability of ERCC1-ERCC4. In total, these observations implicate the GG-NER incision complex and specifically the interaction of USP45 and the disruption of the ERCC1-ERCC4 role in DNA repair as a mechanism of potential importance in MM risk.

Our strategy also identified shared segments overlapping at chr1p36.11 in two Utah pedigrees containing *ARID1A* (Fig. 3 and Table 2) and a genome-wide suggestive segment in a third pedigree harboring another gene in the SWI/SNF complex (*PBRM1*). For the SWI/SNF complex, exome sequencing revealed two rare, potentially deleterious variants in *ARID1A* segregating in two pedigrees. Burden testing provided further evidence for enrichment of variants in *ARID1A* specifically, and in 7 of the 15 genes in the complex. As a component of the SWI/SNF chromatin remodeling complex, ARID1A facilities gene activation by assisting transcription machinery gain access to gene targets [49]. Based on the patterns of mutations in tumor cells, *ARID1A* likely functions as a tumor-suppressor [50]. Members of the SWI/SNF chromatin remodeling complexes are mutated in 20% of malignancies [38], but are extremely intolerant to LoF and missense variation [41] (S5 Table). Blockage of chromatin remodeling may sustain cancer development [39]. Aberrant chromatin remodeling contributes to the pathogenesis of ovarian clear-cell carcinoma [50]. It has previously been shown that *ARID1A* is intolerant to variation (LoF and missense mutations) [28], consistent with its prominent somatic role in multiple tumors [38,50,51], including hematological malignancies [52–54]. These observations implicate the SWI/SNF chromatin remodeling complex, and specifically *ARID1A* in MM risk.

This study has limitations. First, the method is applicable only to extended HRPs that are singularly effective for identifying segregating segments (15 meioses between cases is optimal [16]). The method is not directly applicable to the many smaller family-based resources that have been gathered in the complex trait field and may therefore result in findings from single large pedigrees that are private and difficult to replicate. However, as illustrated in our example, in a collaborative setting containing both extended HRPs and smaller families, the approach can be mutually beneficial. Second, our observation of two borderline genome-wide suggestive overlapping segments at 1p36 led to our identification of *ARID1A* as a potential candidate risk gene and illustrates the potential for discoveries using overlapping subthreshold evidence. However, it raises analytical questions of how to systematically identify such segments. This segment would have been ignored based on strict individual-pedigree thresholds and highlights an important area for further methodological development. Third, as in all family-based genetic studies our method is susceptible to inaccurate pedigree structures and poorly matched control populations. However, relationship and ethnicity checks are standard protocol and mitigate the possibility of error. Finally, this study is observational and cannot describe causation. We have identified two complexes, several genes and specific variants as compelling candidates involved in MM risk, but further functional studies will be required to determine and characterize the mechanisms involved in risk.

In conclusion, we have developed a strategy for gene mapping in complex traits that accounts for heterogeneity within HRPs and formally corrects for multiple testing to allow for statistically rigorous discovery. We applied this strategy to MM, a complex cancer of plasma cells, and identified multiple shared segments containing genes in nucleotide excision repair and SWI/SNF chromatin remodeling. Exome follow-up supported these segments in both the Utah large HRPs and smaller families from other sites. Our study offers a novel technique for HRP gene mapping and demonstrates its utility to narrow the search for risk-variants in complex traits.

## Methods

### SGS Analysis in Utah, Myeloma HRPs

#### HRPs and genotyping

All participants were studied with informed consent under protocols approved by the University of Utah IRB. Using the statewide Utah Cancer Registry (UCR), all living individuals with MM in Utah were invited to participate and peripheral blood was collected for DNA extraction. Participants were linked in the Utah Population Database (UPDB), a unique resource that integrates UCR records with a 5M person genealogy. HRPs were defined as pedigrees containing statistical excess of MM (p<0.05), based on sex and cohort-specific rates in Utah. Eleven of the HRPs identified in the UPDB contained 3-4 MM cases with DNA (total MM cases per pedigree ranged from 4 to 37) with 8-23 meioses between studied MM cases. DNA from the 28 cases was genotyped on the Illumina Omni Express high-density SNP array.

#### Quality control

Only bi-allelic SNPs were considered. Genotypes and individual call-rates were used to ensure high quality data. PLINK was used to remove SNPs with < 95% call rate across individuals [55]. The final SNP set contained 678,447 single nucleotide variants. After SNP removal for low call rates, individuals were removed based on < 90% call rate across the genome, or if they failed the PLINK sex check. One MM case was removed. The QC'ed SNP data were transformed to match strand orientation of the 1000Genomes. PLINK relationship estimates were assessed against pedigree structure from the UPDB to identify any potential issues with pedigree structure. None were found.

#### Probability of sharing a segment

SGS analysis identifies contiguous SNPs that are shared identical-by-state (IBS) by cases in a HRP and assigns an empirical probability of chance ancestral sharing [26]. First, a set of cases in a HRP are defined and all segments of contiguous SNPs shared IBS are identified. All shared segments > 20 SNPs are considered. Lengths shorter than 20 are commonly shared between unrelated individuals. Second, population-based data (here we used CEU and GBR data from the 1000Genomes Project [56]) are used to estimate a graphical model for linkage disequilibrium (LD) [57], providing a probability distribution of chromosome-wide haplotypes in the population. Third, pairs of haplotypes are randomly assigned to pedigree founders according to the haplotype distribution. Founders are individuals whose parents are not specified in the pedigree. For chromosome-wide haplotype simulations the full chromosome LD model is used. Fourth, Mendelian segregation and recombination are simulated to generate genotypes for all pedigree members. The Rutgers genetic map [58] is used for a genetic map for recombination, with interpolation based on physical base pair position for SNPs not represented. Steps two through four create one simulated data set, a random sample from the null hypothesis. This process is repeated hundreds of thousands to millions of times for each subset.

Each shared segment in the real data (step one) is compared to the simulated segments at the precise genomic location. The number of times the null segment equals or encompasses the observed segment is counted and divided by the total number of simulations to generate the empirical nominal p-value for the observed shared segment. The simulations continue until a p-value has been estimated to a required resolution, or until it surpasses a defined significance threshold. To facilitate this in an efficient manner, we follow-up specific segments using marginal distributions from the LD model, established using standard graphical modeling methods [59]. The marginalized LD model encompassing only the region of interest, but capturing relevant LD to accurately simulate genotypes from this region alone. This reduction in markers vastly increases the speed in which simulations are generated. The graphical model estimation, marginalization, and simulation processes are computationally efficient requiring time and storage that is linear with the number of SNPs being considered.

#### Heterogeneity optimization

We systematically perform SGS analysis on each subset of cases in a HRP. If required, the number of subsets can be limited by meioses or subset size. This may be necessary for common traits with large full sets. A lower limit of 10 meioses is a good rule of thumb for reducing the computational burden of subset assessment. At each marker position across the genome, the optimized segment is the one minimizing the p-value across all subsets considered. All segments selected by the optimization procedure, and their respective p-values, comprise the final optimized SGS results.

#### Significance threshold determination

A transformation, *Y* = –*log*_10_(*p*) is performed to the optimized genome-wide SGS p-value vector. The results are fit to a gamma distribution using the MLE method. Y ~Γ(*k*,*σ*) (*k* shape, *σ* rate parameterization). The Theory of Large Deviations has previously been used in pedigree studies to model extreme values in a genomewide genetic setting [27], and it has been shown that for a statistic following a Gaussian distribution, the number of segments where the statistic exceeds a threshold has mean:

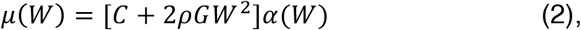

where α(*W*) is the pointwise significance level of exceeding *W*, *C* is the number of chromosomes considered,*ρ* reflects the recombination rate (*ρ* = 1 for general pedigrees), and *G* is genetic length in Morgans. Lander & Kruglyak demonstrated that the same equation extends a statistic following the chi-squared distribution:

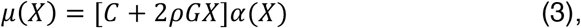

based on the distributional relationship between the chi-squared and Normal distributions *W*^2^ = X. Here, we use the distributional relationship between the gamma and chi-square distributions, our estimated *K* and *σ* gamma parameters, where 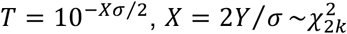, and the genetic length of the genome (matched to that used in the gene-drop) to utilize Eq. 3 and derive μ(*X*) thresholds. Solving for μ(*X*) = 0.05 and μ(*X*) = 1 produced significance and suggestive thresholds, respectively. These thresholds are remarkably stable after a few hundred thousand simulations. For pedigrees with very large numbers of meioses (>50) between the full case-set a larger number of simulations may be required.

#### Software availability

The SGS program is available for download at https://gitlab.com/camplab/sgs and https://gitlab.com/camplab/jps. The main architecture is written in Java. Probability assessments can be multi-threaded, but the largest parallelization gains are achieved by running independent analyses across chromosomes.

### Targeted sequencing

#### Participants

WES data were interrogated in the regions defined by the shared segments of interest. WES data was available on 964 controls [30] and 1,063 MM or MGUS cases including: 28 MM from the 11 Utah HRPs; 70 MM and 46 MGUS from 44 densely clustered families (each containing at least 2 MM or at least 1 MM and 1 MGUS); 186 genetically-enriched MM/MGUS (148 MM and 38 MGUS) including early-onset and MGUS clustering in families; and 733 sporadic MM cases from dbGaP [29]. Of the 44 densely MM/MGUS high-risk families, 25 were ascertained by INSERM, France (36 MM, 38 MGUS), 9 by Mayo Clinic, Minnesota (10 MM, 8 MGUS, 10 unaffected family members), 6 by Memorial Sloan Kettering Cancer Center, New York (14 MM), 3 by International Agency for Research on Cancer, France (8 MM), and 1 by Weill Cornell, New York (2 MM). Most of the families had both MM and MGUS cases (32 families total) and 12 families only had MM cases sequenced. Six families had at least one unaffected relative sequenced. (See S2 Table.) All individuals in the Utah HRPs and the all but three of the densely clustered families were of non-Finish European descent.

#### Joint calling analysis

To perform joint calling of all of the exome sequences, we utilized the calling pipeline developed at the Icahn School of Medicine at Mt. Sinai, based on GATK Best Practices [60]. Briefly, fastq files were aligned to genome build 37 using bwa version 0.7.8, indels were realigned using GATK, duplicates were removed using Picard MarkDuplicates, and base quality scores were recalibrated using GATK. HaplotypeCaller was then used to generate individual GVCF files for each individual, and GenotypeGVCFs was used to generate the final joint calling. The jointly-called VCF was annotated with SNPEff and loaded into a GEMINI (GEnome MINIng) database for ease of querying [61]. Additional functional annotations available in the GEMINI suite include CADD, ANNOVAR, conservation, location, and if the variant was listed in OMIM.

#### Variant prioritization

A GEMINI query was developed to identify variants which were: high or medium impact; AAF < 0.001 in the non-Finnish, European, gnomAD individuals; and within the shared segments of interest. Genes harboring segregating variants in at least two high-risk pedigrees (the discovery pedigree and/or the 44 high-risk pedigrees from collaborating sites) were considered candidate susceptibility genes. These criteria were selected to maintain findings that were unlikely by chance after accounting for both the SGS and sequencing stages of the study. From ExAC exomes, the number of medium/high impact variants with AAF<0.001 per person per gene is 0.0016 [28]. The probability of identifying segregating variants in at least two pedigrees in the same gene can be approximated with a probability from a Binomial(45, 0.0016), which equals 0.0024. To account for the multiple genes in the SGS region, a second probability from Binomial(G, 0.0024) can be used to estimate the probability of observing two segregating variants by chance in G genes. With a threshold of AAF<0.001, the probability of observing at least one gene that harbors 2 variants that segregate in high-risk pedigrees (ø) remains unexpected by chance (<0.05) for up to a reasonable large number of genes (G=20).

#### Joint Assessment of SGS and Sequencing

We assessed the overall rate of expectation for the joint experiments of the SGS and pedigree sequencing findings as π = 11 × µ × ø, where µ is the fully corrected genome-wide rate for the SGS region identified, and ø is the fully corrected probability of the sequencing findings based on the number of genes in the SGS region, as described above.

#### Burden testing

Burden testing was performed on jointly called and processed WES from 1,063 MM/MGUS cases and 964 unaffected controls for the 23 genes in the GG-NER incision complex (including *USP45*) and 15 genes in the SWI/SNF chromatin remodeling complex. The GEMINI software [61] was used to perform a c-alpha test [62] with 1000 permutations. Only variants with AAF < 0.05 and high or moderate predicted impact were included in the analysis.

## Acknowledgments

We thank the DNA Sequencing Core Facility and Genomics Core Facility at the University of Utah, and the computational resources and staff expertise provided by Scientific Computing at the Icahn School of Medicine at Mount Sinai. Data collection was made possible, in part, by the Utah Population Database and the Utah Cancer Registry. We thank the participants and their families who make this research possible.

## Supporting Information

**S1 Fig.**
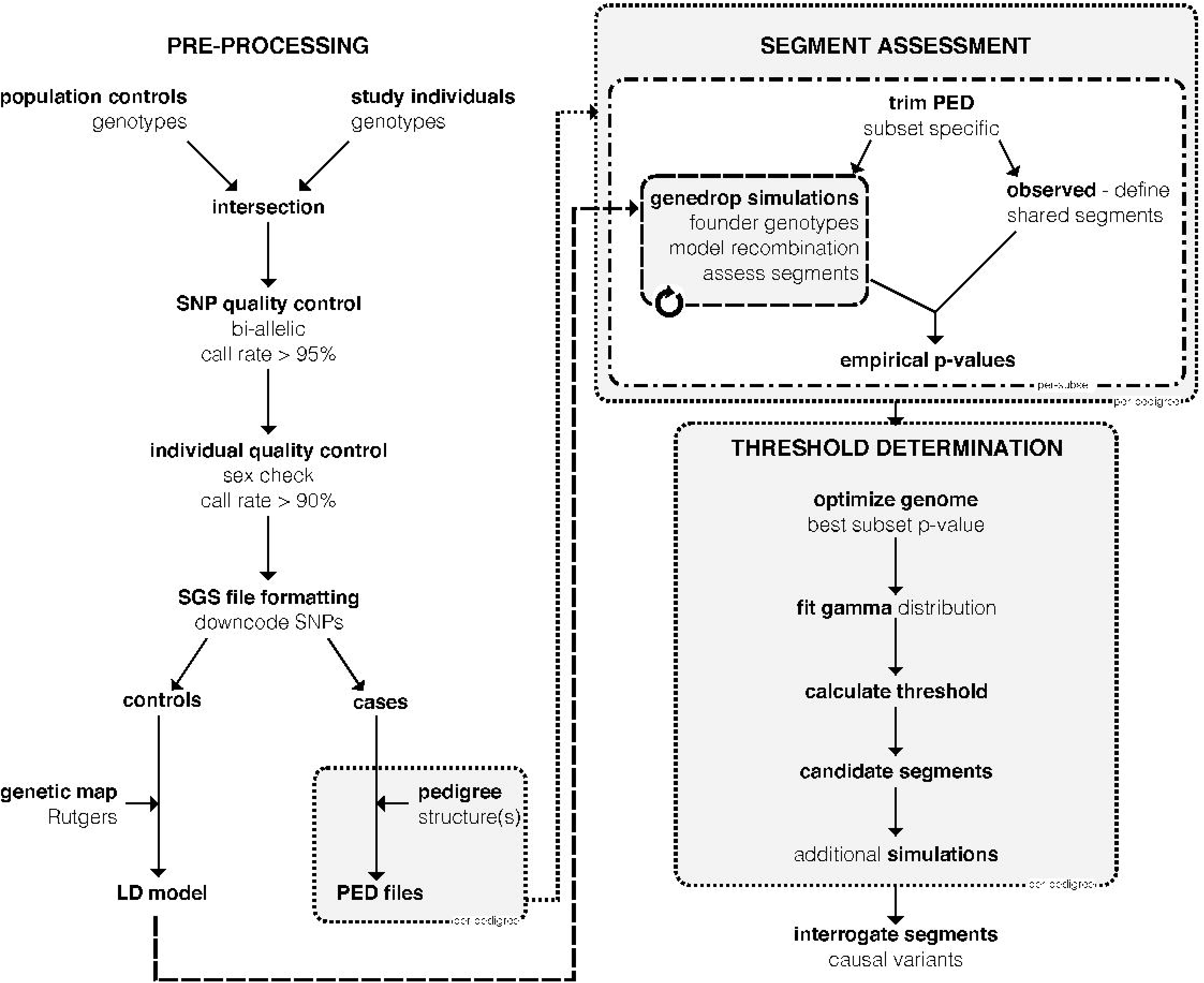
SGS analysis workflow. Overview of the strategy pipeline. Genotypes can be generated from a high-density SNP array, or by extracting SNVs from whole-genome sequencing. CEU and GBR genotypes (unrelated individuals only) from the 1000Genomes Project are generally used as population controls. Dotted boxes represent steps done per-pedigree. Dash-dot boxes represent steps done on all subsets of cases within a pedigree. Dashed box contains step repeated for each simulation. Abbreviations: SNP – single nucleotide polymorphism; SGS – shared genomic segment; LD – linkage disequilibrium; PED – pedigree file (contains relationships and genotypes).

**S2 Fig.**
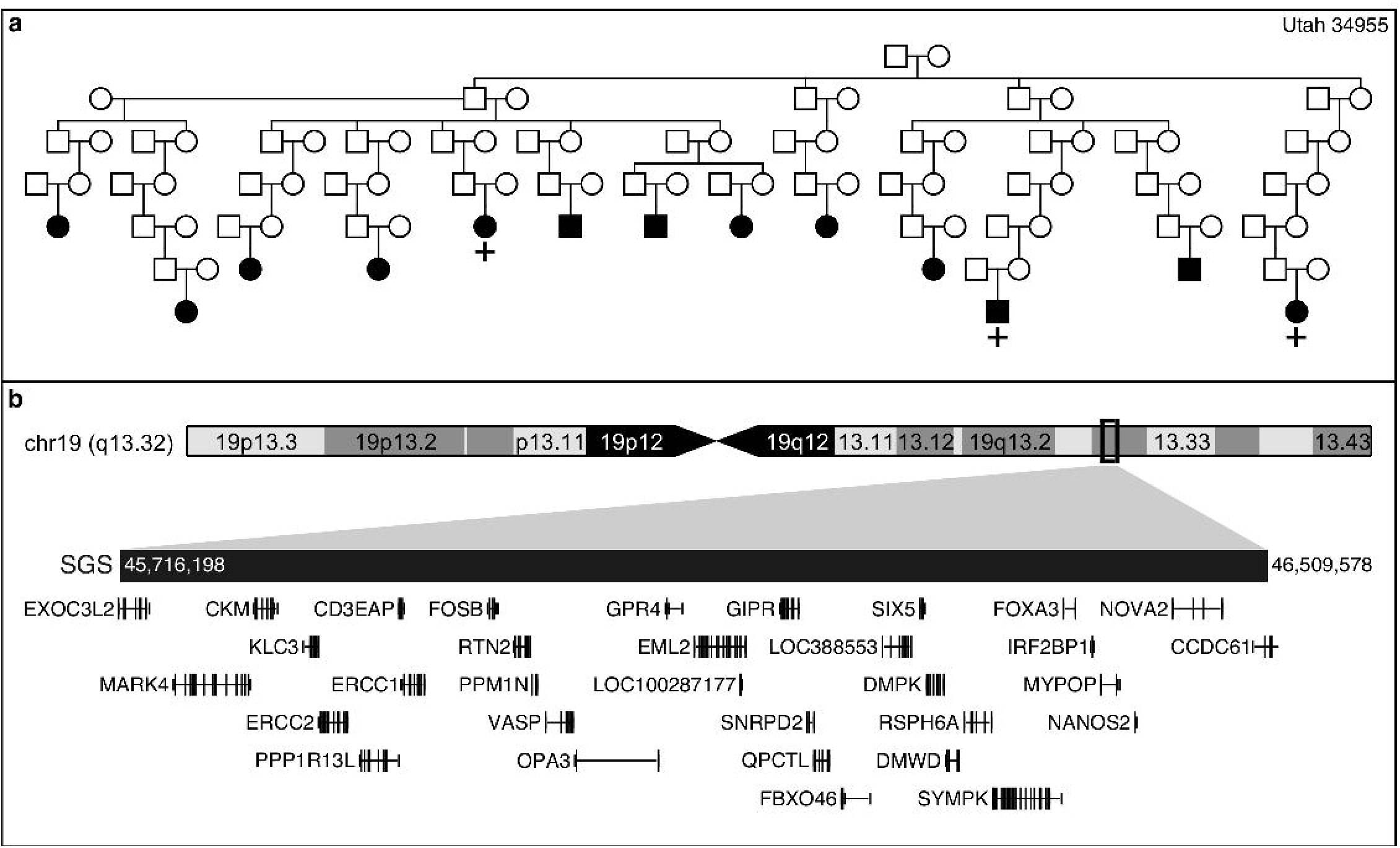
Genome-wide suggestive segment contains *ERCC1*. a) Utah pedigree carrying the genome-wide suggestive SGS at chr19q13.32. + indicates the genotyped MM cases that are SGS carriers, – indicates genotyped and non-carriers, no carrier status indicates not genotyped. b) Genomic region captured by the SGS. *ERCC1* and *ERCC2* are contained.

**S3 Fig.**
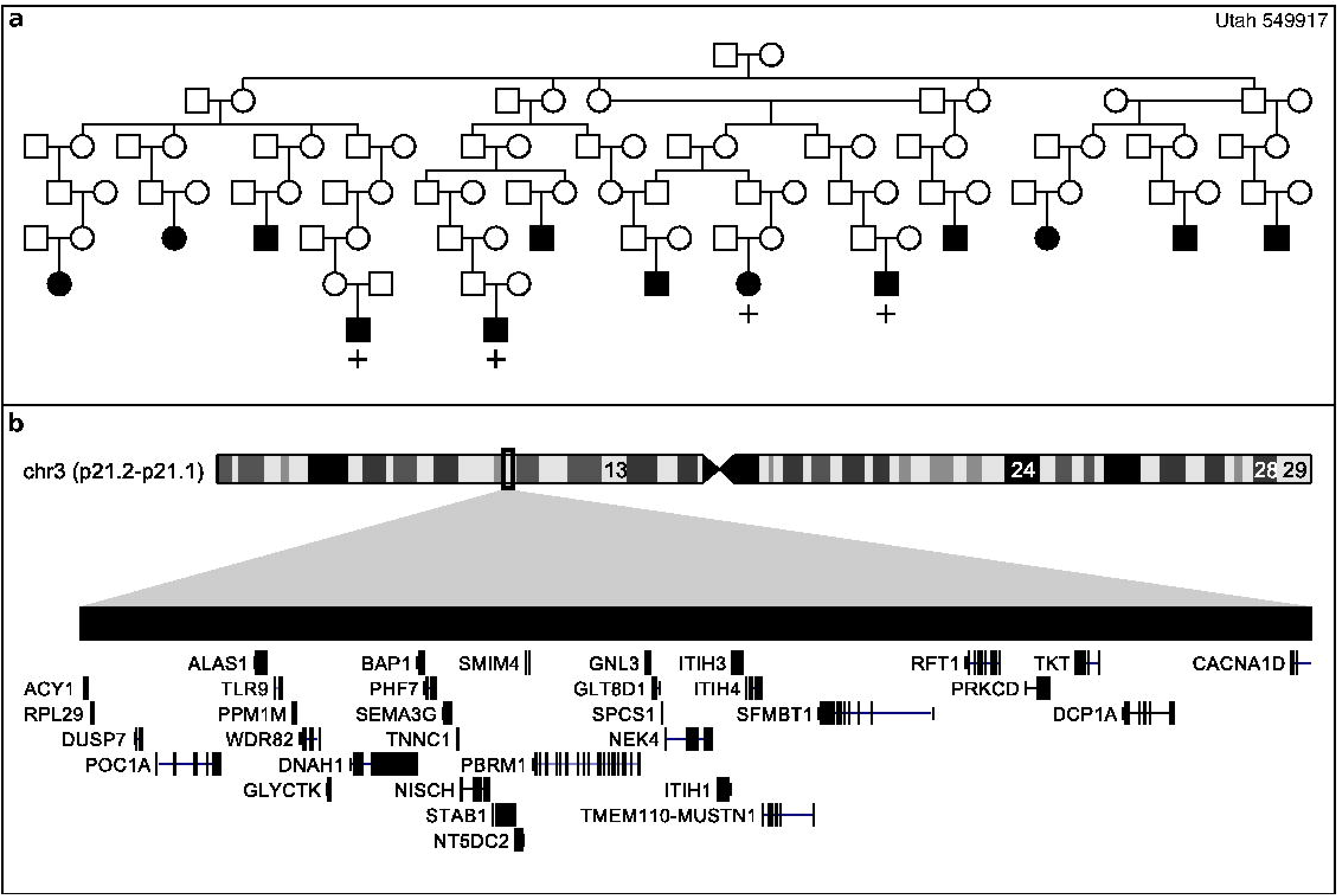
Shared segment containing *PBRM1*. a) Pedigree Utah 549917 carries a genomewide suggestive SGS at chr3p21.2-p21.1. + indicates the genotyped MM cases that are SGS carriers, – indicates genotyped and non-carriers, no carrier status indicates not genotyped. b) Genome region captured by the SGS including *PBRM1*, a component of the SWI/SNF chromatin remodeling complex.

**S1 Table. Genome-wide thresholds and segments.**

**S2 Table. Whole-exome sequenced families.** Total MM, MGUS, and controls in each pedigree and from each site.

**S3 Table. GG-NER Incision Complex genes.** Burden testing results (based on 1063 MM/MGUS cases and 964 unaffected controls), SGS and prioritized SNV results, and tolerance to missense and loss of function variants (based on ExAC population data).

**S4 Table. Evidence for endonuclease regulation of DNA repair.**

**S5 Table. SWI/SNF Complex genes.** Burden testing results (based on 1063 MM/MGUS cases and 964 unaffected controls), SGS and prioritized SNV results, and tolerance to missense and loss of function variants (based on ExAC population data).

**S6 Table. Evidence for SWI/SNF chromatin remodeling.**

